# An apparent lack of synergy between degradative enzymes against *Staphylococcus aureus* biofilms

**DOI:** 10.1101/2023.10.05.561034

**Authors:** Jeremy R. Ellis, Paul A. Rowley

## Abstract

The use of enzymes represents an approach to combat bacterial infections by degrading extracellular biomolecules to disperse *Staphylococcus aureus* biofilms. Commercial enzyme preparations, including cellulase, amylase, pectinase, zymolyase, and pepsin, exhibit concentration-dependent dispersion of *S. aureus* biofilms. Here, we report that low concentrations of these enzymes generally lack synergy when combined or added together sequentially to biofilms. Only the addition of a protease (pepsin) followed by a commercial mixture of degradative enzymes from *Arthrobacter luteus* (zymolyase 20T), demonstrated synergy and was effective at dispersing *S. aureus* biofilms. A more purified mixture of *Arthrobacter luteus* enzymes (zymolyase 100T) showed improved dispersal of *S. aureus* biofilms compared to zymolyase 20T but lacked synergy with pepsin. This study emphasizes the complexity of enzymatic biofilm dispersal and the need for tailored approaches based on the properties of degradative enzymes and biofilm composition.

## Introduction

Biofilms can be constructed by many different microorganisms as an extracellular scaffold that enables surface colonization. The secretion of extracellular polysaccharides and proteins enables surface attachment and the formation of biofilm microcolonies [1,2]. These mature biofilms can be hard to remove and enable the dispersal of planktonic bacterial cells to enable the colonization of new locations. In situations where bacterial colonization is undesirable, biofilms can be a serious challenge and endanger the functionality and longevity of critical infrastructure as well as human health [3–6].

Major components of biofilms are exopolysaccharides (EPS) consisting of homopolysaccharides (i.e cellulose) or heteropolysaccharides (e.g. alginate, Pel, Psl). Digestion of EPS by glycoside hydrolases (GHs) is a strategy that is effective at the removal of bacterial biofilms from surfaces and wounds [7–16]. Many different GHs can degrade EPS constituents of biofilms, such as cellulase, amylase, dispersin B, alginate lyase, and xylosidase, resulting in partial biofilm disruption. In addition to GHs, proteases, and DNases have also been used to disperse biofilms effectively [17–19]. Degradation of biofilms by enzymes allows more effective surface cleaning and increases the efficacy of antibiotics and bacteriophage treatment [16,20–23]. However, not every hydrolytic enzyme is effective at biofilm dispersion, which suggests that the molecular complexity of biofilms and the substrate specificity of enzymes dictates the effectiveness of enzymatic biofilm dispersal [7,11].

Effective biofilm dispersion has been observed with mixtures of GHs and degradative enzymes [7–9,21]. The strategy of using multiple GHs would likely enable the simultaneous disassembly of multiple EPS that would cause a loss of biofilm structural integrity. Targeting different EPS enables the dispersal of complex biofilms that are likely to exist in polymicrobial infections that may be resistant to the action of a single GH. We recently demonstrated that a commercial preparation of cellulase contained a complex mixture of at least two GHs that have different substrate specificities [7]. This “cellulase” mixture degraded the pure substrates carboxymethylcellulose, amylose, and pectin but also efficiently dispersed *Staphylococcus aureus* biofilms. This was in stark contrast to purified recombinant cellulases that were specific in their degradation of carboxymethylcellulose and unable to disperse biofilms. Similarly, the combination of high concentrations of cellulase and amylase mixtures also showed some improvement in biofilm-dispersing activities [21]. Altogether, these data suggest that mixtures of different degradative enzymes can be more effective than purified enzymes. This motivated a systematic approach to understanding how different combinations of degradative enzymes can be used to improve biofilm dispersal. In this study, we investigated the synergistic action of selected GHs, proteases, and DNase to disperse biofilms.

## Results

### A lack of synergy using commercial enzymes to disperse S. aureus biofilms

Previous studies have shown that many different GHs and proteases can degrade biofilms. Therefore, we tested the effectiveness of pectinase, zymolyase, and pepsin in dispersing *S. aureus* biofilm grown on polystyrene (Figure 1A). It was observed that all of these degradative enzymes had concentration-dependent dispersion of biofilms with the most pronounced dispersal at high concentrations of each enzyme (6 mg mL^-1^). Pepsin was found to disperse biofilms at high concentrations despite its low enzymatic activity at pH 7.4. DNAse I was also tested but showed no biofilm degrading activity. To test for potential synergistic activity present between enzymes, we mixed different enzymes at a 1:1 w/w ratio at concentrations that did not exhibit significant biofilm dispersing activity when used alone (DNase I: 50 U/mL and 0.06 mg mL^-1^ for other enzymes). The rationale was that synergy between enzymes would show improved biofilm dispersion compared to the individual enzymes. However, after incubation with *S. aureus* biofilms on polystyrene, there was no observed dispersal with any combination of enzymes (Figure 1B). Enzyme combinations were also mixed at ratios of 1:2, 1:10, and 1:100 with a final enzyme concentration 0.06 mg mL^-1^, but were also ineffective at biofilm dispersal (Figure S1)

**Figure 1.**
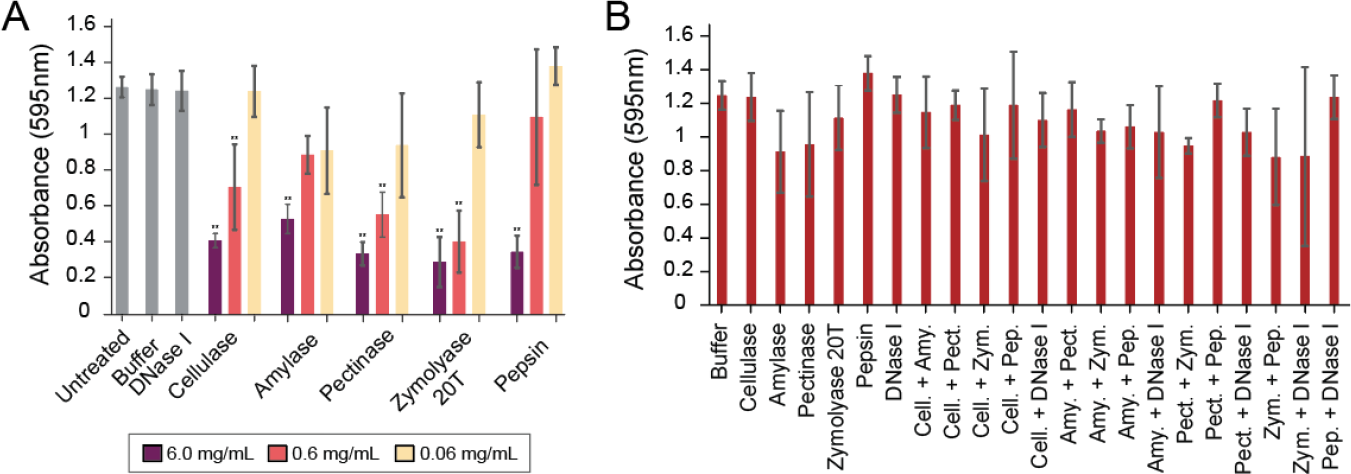
Commercially available enzymes are effective against biofilms at high concentrations but are not synergistic at low concentrations. A. *S. aureus* biofilms grown on polystyrene were treated with three dilutions (6, 0.6, and 0.06 mg mL^-1^) of commercial enzymes (cellulase, amylase, pectinase, zymolyase, and pepsin) and 50 U/mL for DNase I.. B. *S. aureus* biofilms grown on polystyrene were treated with 1:1 mixtures of either 0.06 mg mL^-1^ cellulase (cell.), amylase (amy.), pectinase (pect.), zymolyase 20T (zym.), pepsin (pep.) or 50 U/mL DNase I. (n = 3). After treatment, biofilms were stained with 0.1% crystal violet (** p<0.01, Tukey test).

### The sequential addition of pepsin and zymolyase 20T causes biofilm dispersion

While there was no significant synergy between enzymes when mixed, we were concerned that adding protease could affect the activity of GHs. Therefore, we also assayed whether sequentially adding enzymes at low concentrations would demonstrate synergy. Pepsin and Zymolayse 20T were assayed as they caused some observable biofilm dispersion (Figure 1B), cellulase was included as we had previously characterized the GHs in this enzyme mixture [7]. After one hour of treatment the first enzyme solution was removed and the second enzyme was added and incubated for an additional one hour. Consistent with prior experiments we did not observe any improved biofilm dispersing activities for most combinations of enzymes. However, the altered treatment regime did reveal that adding pepsin followed by zymolase 20T caused a significant dispersion of *S. aureus* biofilms compared to adding either of the enzymes alone (Figure 2A). Importantly, reversing the order of addition so that zymolase 20T was added first, followed by pepsin, did not result in biofilm dispersal. A multispecies biofilm containing *Pseudomonas aeruginosa* and *S. aureus* was also used to test synergy, but no treatment resulted in biofilm dispersion (Figure 2B) [24].

**Figure 2.**
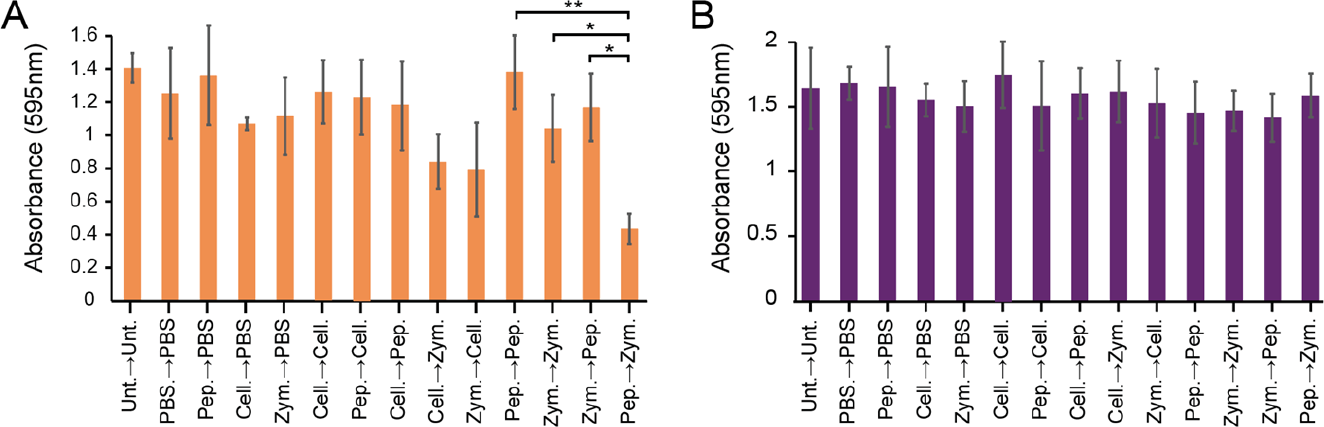
Sequential treatment of pepsin followed by zymolyase 20T leads to *S. aureus* biofilm dispersal. A. *S. aureus* and B. multi-species (*S. aureus*, and *P. aeruginosa*) biofilms grown on polystyrene were treated sequentially with the indicated enzymes for one hour per treatment phase. Different sequential combinations of cellulase (cell.), zymolyase 20T (zym.), and pepsin (pep.)) were added to a final enzyme concentration of 0.06 mg mL^-1^. Unt, untreated; PBS, phosphate-buffered saline. (* p<0.05, ** p<0.01, Tukey test) (n = 3).

### Zymolyase 100T is highly effective at dispersing biofilms but lacks synergy with pepsin

Zymolyase 20T is a commercially available preparation of multiple enzymes precipitated by ammonium sulfate from cultures of the bacteria *Arthrobacter luteus*. This preparation is typically used to digest cell wall components of multiple different species of fungi [25]. Many different enzymatic activities are present in these commercial preparations, including ß-glucanase, protease, and mannanase. Enzymes associated with ß-glucanase and protease activities have been successfully separated by additional steps of protein purification [26]. Indeed, zymolyase 100T is a preparation derived from 20T that has been purified by ß-1,3-glucan affinity chromatography to enrich the ß-glucanase activity [27]. We find that the 100T preparation appears to have a similar composition of proteins compared to 20T by SDS-PAGE gel electrophoresis (Figure S2), but is significantly more effective at dispersing biofilms of *S. aureus* (Figure 3A). Specifically, 100T causes significant biofilm dispersal at a concentration of 6e-4 ug mL^-1^ compared to 0.6 mg mL^-1^ for 20T, which is likely explained by the increased concentration of ß-glucanase in the preparation due to affinity purification. However, the sequential addition of pepsin followed by 100T failed to elicit synergy, as was observed with 20T (Figure 3B).

**Figure 3.**
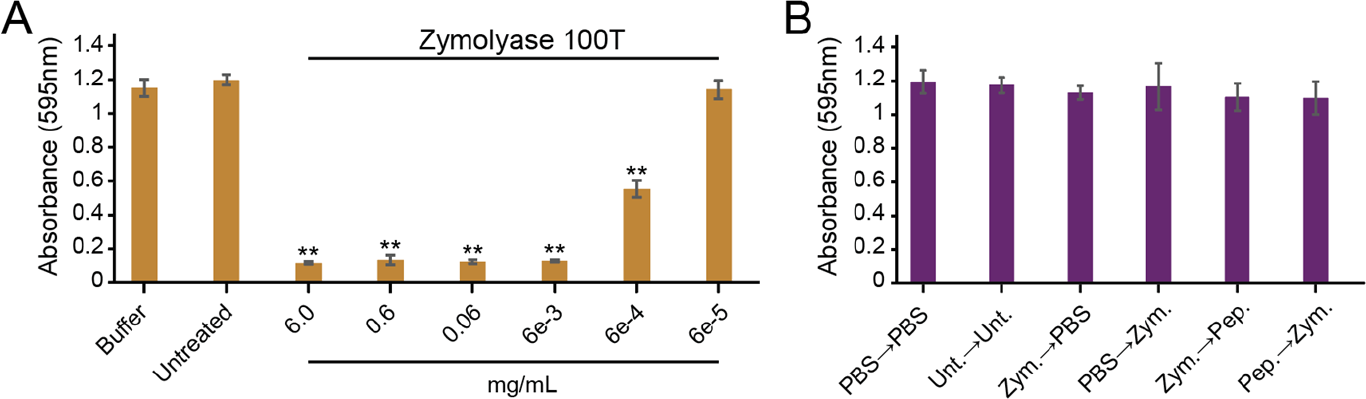
Zymolyase 100T is highly effective at dispersing *S. aureus* biofilms but lacks synergy with pepsin. A. *S. aureus* biofilms grown on polystyrene were treated with zymolyase 100T at different concentrations for one hour. B. *S. aureus* biofilms grown on polystyrene were treated sequentially with zymolyase 100T (Zym.; 6e-5 ug mL^-1^) and pepsin (Pep.; 0.06 mg mL^-1^). Biofilms were stained with 0.1% crystal violet, and the absorbance was measured at 595 nm (* p<0.05, ** p<0.01, Tukey test) (n = 3).

## Discussion and Conclusion

GHs have been shown to efficiently degrade bacterial biofilms, which holds future promise for combatting recalcitrant bacterial infections that are resistant to antibiotics. The motivation behind the current study was to identify combinations of enzymes to disperse biofilms more efficiently when compared to single enzymes. As biofilms are often composed of many different polymers, including proteins, carbohydrates, and nucleic acids, it seemed reasonable that combining biofilm-degrading enzymes would lead to an enhancement of dispersal. This would be analogous to approaches to the deconstruction of complex plant cell wall carbohydrates for biofuel production [28]. However, although we found that many enzymes were effective at the dispersal of biofilms at high concentrations, no pair of enzymes demonstrated synergy when combined at concentrations that would not cause dispersal. However, the sequential addition of pepsin followed by zymolyase 20T was unique in its ability to cause biofilm dispersion, whereas the reverse order of addition showed no effect. This result is similar to studies of plant cell wall digestion experiments that have determined that carbohydrate-binding modules can potentiate enzymatic degradation [29]. Specifically, the binding of these carbohydrate-binding modules is thought to unmask carbohydrates of the plant cell wall that are susceptible to degradation by enzymes. Similarly, the action of non-hydrolytic accessory enzymes on cellulose can also enable more efficient depolymerization by cellulases [28]. These requirements highlight the importance of order-of-addition for the deconstruction of complex organic substrates that could apply to the dispersal of biofilms. We draw an analogy between plant cell walls and bacteria biofilms as both are known to contain ordered structures of carbohydrates [30,31]. In the current study, it is feasible that unmasking carbohydrates by the action of pepsin, either by proteolysis or the binding of biofilms, could allow the hydrolysis of previously masked carbohydrates upon the addition of 20T. *S. aureus* is known to encode extracellular proteases that are thought to be required for biofilm formation and remodeling [32]. Specifically, *S. aureus* proteases can cleave the extracellular *S. aureus* Biofilm Associated Protein (BAP), which is essential for adhesion and biofilm development [33]. This is only one of many proteins that are integral to bacterial biofilms and could be targeted by proteolysis to alter biofilm structure (such as [34–36]). However, we find that combining different degradative enzymes mostly does not enhance biofilm dispersion, despite prior observations that mixtures of GHs and other biologically active molecules can have an additive effect on dispersion [7–9,16].

Our previous work has shown that commercial GHs can contain a complex mixture of different proteins with varying hydrolytic activities [7]. This adds uncertainty to the interpretation of the effects of degradative enzymes on biofilms, as it is difficult to pinpoint the relevant enzymatic activities required for dispersion. Like other commercial preparations of GHs, zymolyase 20T is a crude mixture of different enzymes extracted by ammonium sulfate precipitation. Specifically, the manufacturer’s specification for 20T detected the presence of significant ß-glucanase, mannanase, and protease activities. In the current study, there was a clear difference in the composition of zymolyase 20T and 100T that affected biofilm dispersing synergy with pepsin, even if there was not a dramatic difference in the overall protein composition of the mixture. The vital component that enabled synergy with pepsin was likely removed after affinity purification of 100T using ß-1,3-glucan [27]. Further analysis of the composition of 20T compared to 100T would be required to determine the molecule in 20T responsible for synergy. It is also important to recognize that the purity of the pepsin used in the current study was not tested and could contain other biologically active compounds with the potential to influence synergy. Pepsin and 20T were also unable to disperse a two-species biofilm, demonstrating that synergy was specific to the composition of *S. aureus* biofilms that was lost upon the addition of *P. aeruginosa*. It is well understood that biofilms vary in their composition, which would likely influence the effectiveness of enzymatic biofilm dispersion, adding an additional layer of complexity to the use of GHs for biofilm dispersal. This highlights the need to further investigate the role of synergy in biofilm dispersal with more diverse combinations of purified GHs and other degradative enzymes.

## Materials and Methods

### Propagation of microorganisms

The bacterial strains *S. aureus* SA31 and *P. aeruginosa* PAO1 were used for biofilm studies and maintained by growing in tryptic soy broth (TSB) and Luria broth (LB), respectively.

### Enzymes used for biofilm dispersion

Enzymes used in this study were sourced from various companies; cellulase (*Aspergillus niger*) MP Biomedicals, catalog number 150583; amylase (*Bacillus sp*.), MP Biomedicals, catalog number 100447; pectinase (*Rhizopus sp*.), Sigma-Aldrich, catalog number P2401; zymolyase 20T (*Arthrobacter luteus*), Amsbio, catalog number 120491-1; zymolyase 100T (*Arthrobacter luteus*), United States Biological, catalog number Z1004; pepsin (Porcine gastric mucosa), Sigma-Aldrich, catalog number P7000; DNase I, Zymo Research, catalog number E1010. All enzymes were suspended in phosphate-buffered saline at pH 7.4 to the desired concentration.

### Polystyrene biofilm model

The *in vitro* polystyrene biofilm model used to measure the effectiveness of GHs at dispersion is described in our prior manuscript [7]. For the sequential treatment of biofilms with enzymes, the first enzyme treatment was allowed to bathe biofilms for 1 hour, after which the liquid was carefully aspirated. The second enzyme treatment was added for an additional hour. Crystal violet staining and subsequent quantification occurred only after the second treatment course was completed. The multispecies biofilm with *S. aureus* SA31 and *P. aeruginosa* PAO1 was constructed as described previously [24], but with the following modifications: after the mixed bacterial suspension was inoculated in 96-well round-bottomed polystyrene plates, biofilms were allowed to grow for 48 h at 37°C. Supernatants were aspirated and the biofilms were treated as described in our prior manuscript for single-species *S. aureus* biofilms.

### Analysis of Zymolyase 20T and 100T by SDS-PAGE

10 µg of 20T and 100T zymolyase were run on a 10% SDS-PAGE gel and subsequently visualized with Coomassie staining.

## Acknowledgments

We would like to acknowledge the mentorship and support of Professor James J. Bull at the University of Idaho.

## Supplement

**Figure S1.**
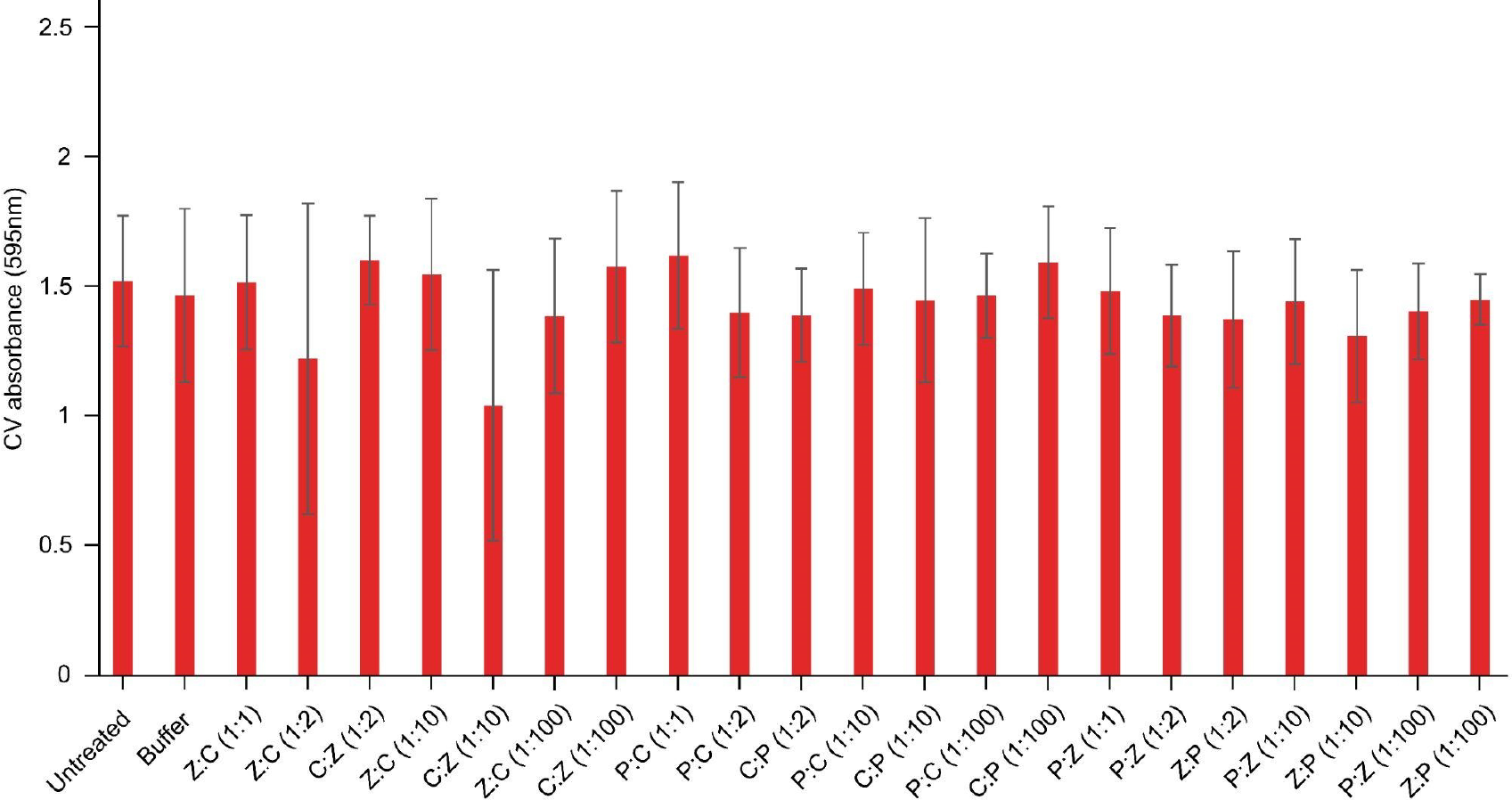
No synergy between different ratios of enzymes at a low concentration. *S. aureus* biofilms grown on polystyrene were treated with mixed enzyme combinations for one hour. Enzyme combinations were mixed at ratios of 1:1, 1:2, 1:10, and 1:100 with total enzyme concentration at 0.06 mg mL^-1^. Wells were stained with 0.1% crystal violet, and the absorbance was measured at 595 nm (P = pepsin, Z = zymolyase 20T, C = cellulase) (n = 3).

**Figure S2.**
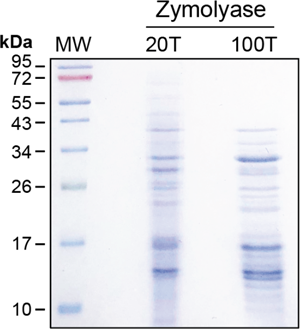
Analysis of Zymolyase 20T and 100T by SDS PAGE. 10 ug of zymolyase 20T and 100T were analyzed by 10% SDS PAGE. The molecular weight marker used in lane 1 was the Color Prestained Protein Standard, Broad Range (BioRad).

